# Altered expression of a unique set of genes reveals complex etiology of Schizophrenia

**DOI:** 10.1101/131623

**Authors:** Ashutosh Kumar, Himanshu Narayan Singh, Vikas Pareek, Khursheed Raza, Pavan Kumar, Muneeb A. Faiq, Sankat Mochan, Subrahamanyam Dantham, Ashish Datt Upadhyaya

## Abstract

**Purpose:** The etiology of schizophrenia is extensively debated, and multiple factors have been contended to be involved. A panoramic view of the contributing factors in a genome-wide study can be an effective strategy to provide a comprehensive understanding of its causality.

**Materials and Methods:** GSE53987 dataset downloaded from GEO-database, which comprised mRNA expression data of post-mortem brain tissue across three regions from control and age-matched subjects of schizophrenia (N= Hippocampus (HIP): C-15, T-18, Prefrontal cortex (PFC): C-15, T-19, Associative striatum (STR): C-18, T-18). Bio-conductor-affy-package used to compute mRNA expression, and further t-test applied to investigate differential gene expression. The analysis of the derived genes performed using PANTHER Classification System and NCBI database.

**Results:** A set of 40 genes showed significantly altered (p<0.01) expression across all three brain regions. The analyses unraveled genes implicated in biological processes and events, and molecular pathways relating basic neuronal functions.

**Conclusions:** The deviant expression of genes maintaining basic cell machinery explains compromised neuronal processing in SCZ.

**Abbreviations:** Schizophrenia (SCZ), Hippocampus (HIP), Associative striatum (STR), Prefrontal cortex (PFC)

## Introduction

The etiology of Schizophrenia (SCZ) is extensively debated [1,2]. An uncertainty of the etiology has greatly impeded the treatment of the disease, and neither of the therapeutic approaches [1] is proving much helpful in halting its progression. Many candidate genes have been reported [3] but none of them actually got validated in population-based studies for persistent association [4,5]. A disease signature derived from genome-wide expression patterns in affected brain regions was highly desirable that would not only help to reach to accurate diagnosis of the disease but also in developing optimal therapeutic approaches aimed at maximum relief of the patients.

### SCZ pathology may be reflected in expression analysis of neural genes

SCZ has been noted to cause significant architectural changes in many brain regions, the hippocampal, prefrontal cortex, and basal nuclei regions have been chief among them [6-8]. The architectural changes in the brain varied from the changes in the total size and volume of specific brain regions [9,10] to neural connection between different brain regions [11-13], pruning of dendritic spines [14] and synapses [15,16], synaptic dysfunction [17], and also functional changes as oscillatory coupling [18] and neuronal firing patterns [19]. Each SCZ brain may have a few or more of such architectural defects. The recent study by Sekar et al (2016) in mice models reported a variant allelic form of complement C4A (which is involved into the pruning of synapses) may cause excessive synaptic pruning in developing neural circuits and may put the individuals at risk of developing SCZ [15].

How the neural architectural changes are instructed by the changes in the neural genes has also been shown by some recent studies. Piskorowski et al (2016) have shown in the mouse model that deletion of 22q11 locus may involve the genes making synaptic proteins and that may produce SCZ like symptoms [20]. Fromer and colleagues (2016) have identified over 100 of genetic loci harbouring SCZ associated variants which together involve scores of genes supporting polygenic etiology of SCZ, and also altering the expression or knock down of some of such genes in animal or human stem cell models has shown to compromise neural functions effectively [21].

### Genetic basis of SCZ: much is now known but connecting mechanism is missing

SCZ gives a life time risk of ~1 % and shows high heritability (~ 69-81%) [22-24]. The SCZ heritability is derived from CNVs, SNPs, de novo mutations, and structural modifications at gene promoter regions without involving gene sequences as have been revealed in the genome-wide studies [25-28]. Plenty of CNVs and SNP variants have been reported until [29] yet but none of these appear to be present as a constant association, and also none of them ensures a causal association or contributing alone significantly to the genetic liability for the disease. Emerging evidence suggest the genetic etiology of SCZ may be deriving from accumulative effect of all such gene structure changes [21,30] which plausibly act through influencing expression of neuronal genes by altering gene promoter regions [21,31]. These factors together may implicate thousands of the genes, an indication for the polygenic etiology of the SCZ [21,30,32,33].

Expression derangement of the genes also evidenced to arise of the gene-environment interactions during foetal development and in the lifetime of the individuals [25,34,35] involving the mechanisms as de novo mutations, and epigenetic modifications at the gene promoter sites [27,33,36].

The biological contribution to the disease etiology is ascertained from the adoption studies which showed that offspring of the diseased mothers, although adopted by normal families, bear high risk for developing the disease [37]. Conversely, an essential environmental contribution to the disease etiology is indicated by the findings in the twin-based studies that the siblings of the monozygotic twins, who although share almost same genome, but differ in heritability for SCZ [38]. A gene-environmental interaction necessary for the disease etiology was further indicated by the observation in the adoption studies that siblings of the mothers with schizophrenia showed more prevalence of the disease in harsher rearing conditions in adopter families in comparison to the controls whose original parents had no disease [37].

Prenatal and perinatal environments have been also evidenced to influence initiation of the disease in the adult [39]. Prenatal maternal infections and psychological stress, and also obstetric conditions have been found to put permanent influence on the foetal brain and are considered risk factors for schizophrenia [40-42]. Evidence supports the view that insults to the developing brain get hardwired which creates susceptibility for developing SCZ in adulthood if faced with stressful life conditions [43-45].

Furthermore, even the maternal depression or severe stress in perinatal and/or in childhood period have been found to raise chances of SCZ in the offspring during adulthood [46,47].

The ‘two-hit’ hypothesis for etiogenesis of SCZ [43,45,48] has got wide acceptance among scholars and now a ‘multiple hit’ hypothesis is being suggested by some authors [49]. Literature evidence suggest that CNVs and SNPs, specific mutations and also intergenerational epigenetic influences create a primarily genetically susceptible brain which if gets further hit by adverse environmental conditions during adulthood, may be leading to SCZ [50-53]. Similarly, a significant environmental insult during development may also prime the brain for developing SCZ with further environmental hits [44].

The challenging environmental conditions during pre and perinatal period [39,54,55], childhood rearing up, and adolescence which found associated with increased risk of SCZ, all may be involved in SCZ etiogenesis by influencing the expression of neuronal genes [31].

Social environmental conditions during adulthood like unemployment, urban living, geographical migration, and prolonged war have been also reported to be associated with increased chances of getting SCZ [56,57]. The mediating mechanism for social environment induced SCZ seems to be chronic psychological stress acting on the neural genes mediated by different complex biological methods as intergenic interactions (epistasis), epigenetic reprogramming, or microRNA [58-59] or splicing quantitative trait loci (sQTLs) mediated regulations [60]. The psychological stress conditions as early life bereavement, social defeats may also be a precondition for SCZ [46,61,62].

Whatever be the organizing mechanism, the resultant up and down regulation of the neuronal genes may be the chief etiological mechanism in SCZ [31]. Plausibly, the dysregulation of the neuronal genes, especially which are involved in maintaining basic cell architecture and machinery, may compromise the information processing in neurons in affected brain regions which manifests as disorganized and deficient behaviour evident in SCZ [63,64].

### Ontological analysis of altered neural genes may reflect SCZ etiology

Based on the mounting body of evidence we discussed above, we hypothesized that the altered expression of neuronal genes may be the connecting link between all genetic and environmental factors involved in the etiogenesis of SCZ; hence an ontological analysis of the genes showing altered expression in various brain regions may unravel the components of the complex etiology of SCZ.

## Materials & Methods

### Data Resources

The mRNA expression data were retrieved from the GEO (Genome Expression Omnibus, GSE53987) (http://www.ncbi.nlm.nih.gov/geo/), a public repository for high-throughput microarray. The RNA was originally isolated from post-mortem brain tissue across three specific regions (Hippocampus (HIP), Prefrontal cortex (PFC): Brodmann Area 46, and Associative striatum (STR)) of control {N=18 (HIP), 19 (PFC), 18(STR)} and age-matched subjects with schizophrenia {N=15 (HIP), 15 (PFC), 18 (STR)}. Equal numbers of male and female (except for odd number samples) diagnosed SCZ cases and controls of adult age (range 22-68 years). The controls were matched for the age and sex with cases, and were free of any neurological or psychiatric illness during their life course. Randomized sampling of the cases and controls were applied to keep away the selection biases.

### Data retrieval and analysis

The RNA was isolated from HIP, PFC (Brodmann Area 46), and associative STR and hybridized to U133_Plus2 Affymetrix chips for m-RNA expression study. Expression analysis of mRNA was done by using “affy” package (http://www.bioconductor.org/packages/release/bioc/html/affy.html), which was deposited at Bioconductor and developed in R statistical software program and scripting language. It used three steps to calculate the expression intensities: (i) background correction; (ii) normalization (data were normalized by RMA, subjected to pair wise comparison followed by Benjamini and Hochberg False Discovery rate correction (FDR)), and (iii) expression calculation. After calculation of mRNA expression intensity a simple unpaired two tailed t-test (significance set at p ≤0.01) was applied to the data to filter out the set of genes expressed significantly in all three brain regions.

To categorize the derived significantly altered genes on the basis of their involvement in molecular functions, molecular pathways, and biological events, PANTHER (Protein ANalysis THrough Evolutionary Relationships) Classification System (http://www.pantherdb.org/) and NCBI gene database (http://www.ncbi.nlm.nih.gov/gene/) were exploited.

## Results

A set of 40 genes (Protein coding-38; RNA-gene-2) was identified showing statistically significant (p ≤0.01) altered mRNA expression in schizophrenic patients in the all three brain regions studied (Table 1). Interestingly, it was observed that most of the genes were down-regulated in all three brain regions (32/40). Also, the same genes in all three brain regions have shown the similar direction of expression changes.

**Table 1.**
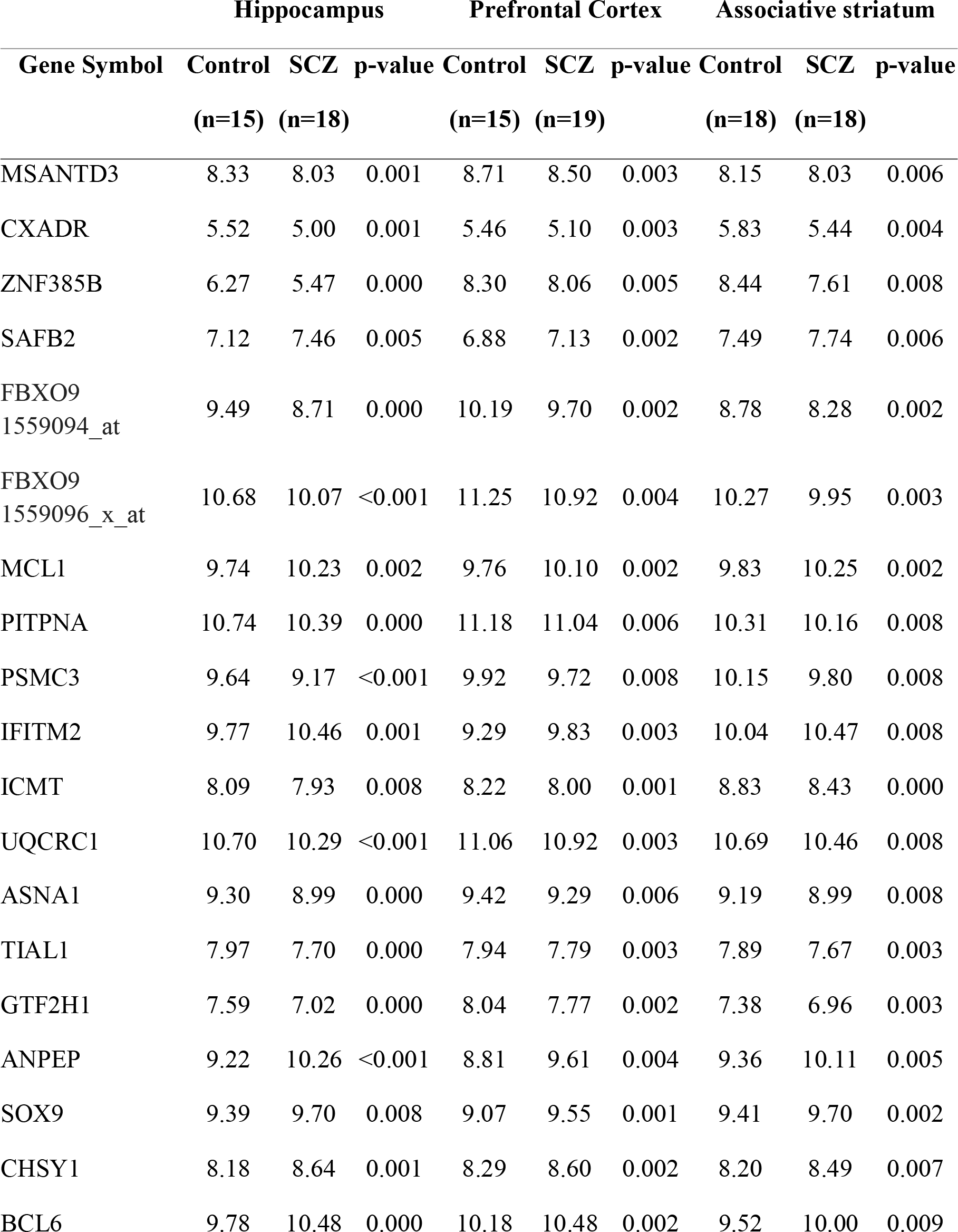
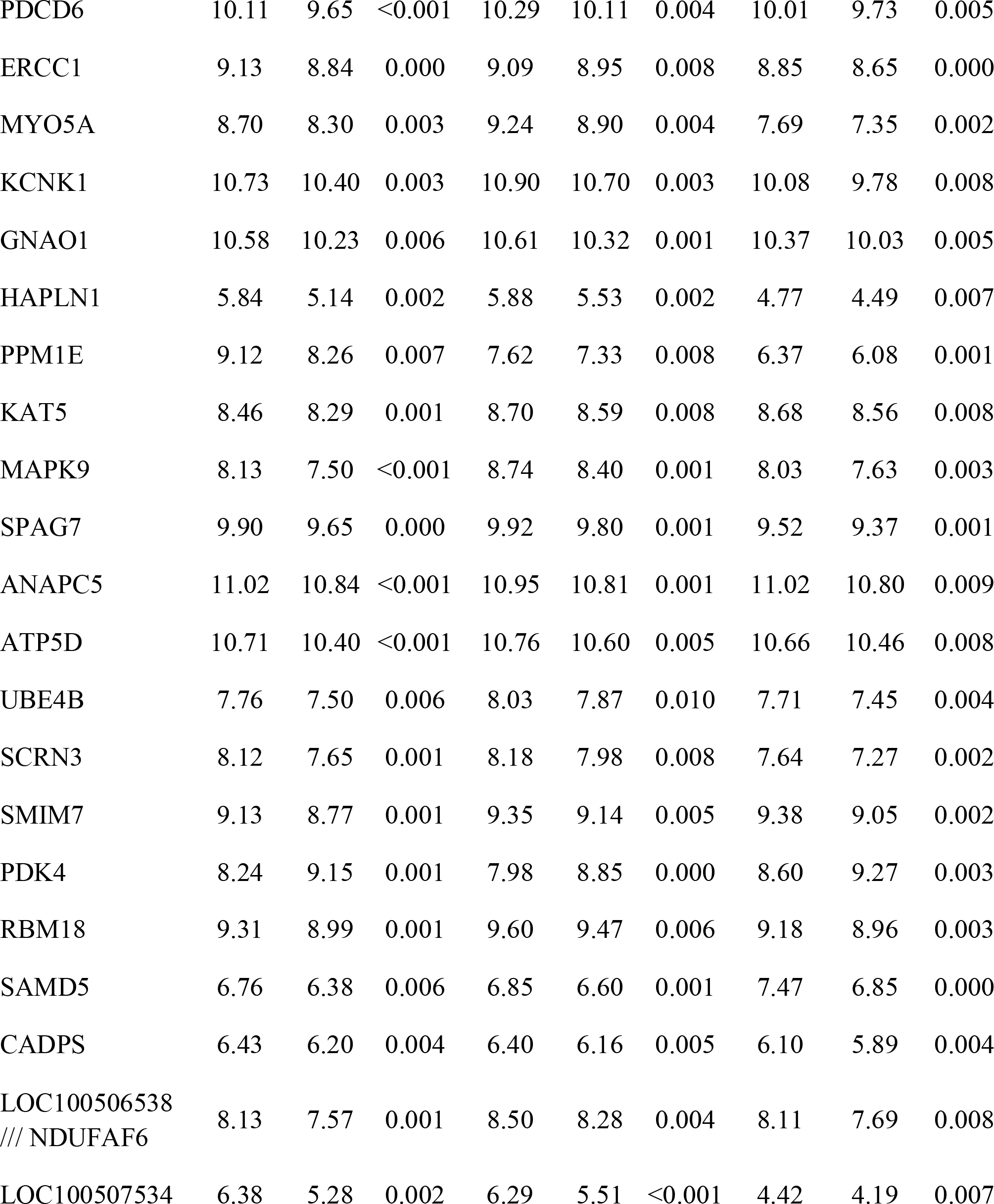
Genome wide m-RNA expression (statistical significance set at p ≤0.01) in three brain regions of schizophrenic patients and healthy controls (data represented as mean).

|  | Hippocampus |  |  | Prefrontal Cortex |  |  | Associative striatum |  |  |
| --- | --- | --- | --- | --- | --- | --- | --- | --- | --- |
| Gene Symbol | Control | SCZ | p-value | Control | SCZ | p-value | Control | SCZ | p-value |
|  | (n=15) | (n=18) |  | (n=15) | (n=19) |  | (n=18) | (n=18) |  |
| MSANTD3 | 8.33 | 8.03 | 0.001 | 8.71 | 8.50 | 0.003 | 8.15 | 8.03 | 0.006 |
| CXADR | 5.52 | 5.00 | 0.001 | 5.46 | 5.10 | 0.003 | 5.83 | 5.44 | 0.004 |
| ZNF385B | 6.27 | 5.47 | 0.000 | 8.30 | 8.06 | 0.005 | 8.44 | 7.61 | 0.008 |
| SAFB2 | 7.12 | 7.46 | 0.005 | 6.88 | 7.13 | 0.002 | 7.49 | 7.74 | 0.006 |
| FBXO9<br>1559094_at | 9.49 | 8.71 | 0.000 | 10.19 | 9.70 | 0.002 | 8.78 | 8.28 | 0.002 |
| FBXO9<br>1559096_x_at | 10.68 | 10.07 | <0.001 | 11.25 | 10.92 | 0.004 | 10.27 | 9.95 | 0.003 |
| MCL1 | 9.74 | 10.23 | 0.002 | 9.76 | 10.10 | 0.002 | 9.83 | 10.25 | 0.002 |
| PITPNA | 10.74 | 10.39 | 0.000 | 11.18 | 11.04 | 0.006 | 10.31 | 10.16 | 0.008 |
| PSMC3 | 9.64 | 9.17 | <0.001 | 9.92 | 9.72 | 0.008 | 10.15 | 9.80 | 0.008 |
| IFITM2 | 9.77 | 10.46 | 0.001 | 9.29 | 9.83 | 0.003 | 10.04 | 10.47 | 0.008 |
| ICMT | 8.09 | 7.93 | 0.008 | 8.22 | 8.00 | 0.001 | 8.83 | 8.43 | 0.000 |
| UQCRC1 | 10.70 | 10.29 | <0.001 | 11.06 | 10.92 | 0.003 | 10.69 | 10.46 | 0.008 |
| ASNA1 | 9.30 | 8.99 | 0.000 | 9.42 | 9.29 | 0.006 | 9.19 | 8.99 | 0.008 |
| TIAL1 | 7.97 | 7.70 | 0.000 | 7.94 | 7.79 | 0.003 | 7.89 | 7.67 | 0.003 |
| GTF2H1 | 7.59 | 7.02 | 0.000 | 8.04 | 7.77 | 0.002 | 7.38 | 6.96 | 0.003 |
| ANPEP | 9.22 | 10.26 | <0.001 | 8.81 | 9.61 | 0.004 | 9.36 | 10.11 | 0.005 |
| SOX9 | 9.39 | 9.70 | 0.008 | 9.07 | 9.55 | 0.001 | 9.41 | 9.70 | 0.002 |
| CHSY1 | 8.18 | 8.64 | 0.001 | 8.29 | 8.60 | 0.002 | 8.20 | 8.49 | 0.007 |
| BCL6 | 9.78 | 10.48 | 0.000 | 10.18 | 10.48 | 0.002 | 9.52 | 10.00 | 0.009 |
| PDCD6 | 10.11 | 9.65 | <0.001 | 10.29 | 10.11 | 0.004 | 10.01 | 9.73 | 0.005 |
| ERCC1 | 9.13 | 8.84 | 0.000 | 9.09 | 8.95 | 0.008 | 8.85 | 8.65 | 0.000 |
| MYO5A | 8.70 | 8.30 | 0.003 | 9.24 | 8.90 | 0.004 | 7.69 | 7.35 | 0.002 |
| KCNK1 | 10.73 | 10.40 | 0.003 | 10.90 | 10.70 | 0.003 | 10.08 | 9.78 | 0.008 |
| GNAO1 | 10.58 | 10.23 | 0.006 | 10.61 | 10.32 | 0.001 | 10.37 | 10.03 | 0.005 |
| HAPLN1 | 5.84 | 5.14 | 0.002 | 5.88 | 5.53 | 0.002 | 4.77 | 4.49 | 0.007 |
| PPM1E | 9.12 | 8.26 | 0.007 | 7.62 | 7.33 | 0.008 | 6.37 | 6.08 | 0.001 |
| KAT5 | 8.46 | 8.29 | 0.001 | 8.70 | 8.59 | 0.008 | 8.68 | 8.56 | 0.008 |
| MAPK9 | 8.13 | 7.50 | <0.001 | 8.74 | 8.40 | 0.001 | 8.03 | 7.63 | 0.003 |
| SPAG7 | 9.90 | 9.65 | 0.000 | 9.92 | 9.80 | 0.001 | 9.52 | 9.37 | 0.001 |
| ANAPC5 | 11.02 | 10.84 | <0.001 | 10.95 | 10.81 | 0.001 | 11.02 | 10.80 | 0.009 |
| ATP5D | 10.71 | 10.40 | <0.001 | 10.76 | 10.60 | 0.005 | 10.66 | 10.46 | 0.008 |
| UBE4B | 7.76 | 7.50 | 0.006 | 8.03 | 7.87 | 0.010 | 7.71 | 7.45 | 0.004 |
| SCRN3 | 8.12 | 7.65 | 0.001 | 8.18 | 7.98 | 0.008 | 7.64 | 7.27 | 0.002 |
| SMIM7 | 9.13 | 8.77 | 0.001 | 9.35 | 9.14 | 0.005 | 9.38 | 9.05 | 0.002 |
| PDK4 | 8.24 | 9.15 | 0.001 | 7.98 | 8.85 | 0.000 | 8.60 | 9.27 | 0.003 |
| RBM18 | 9.31 | 8.99 | 0.001 | 9.60 | 9.47 | 0.006 | 9.18 | 8.96 | 0.003 |
| SAMD5 | 6.76 | 6.38 | 0.006 | 6.85 | 6.60 | 0.001 | 7.47 | 6.85 | 0.000 |
| CADPS | 6.43 | 6.20 | 0.004 | 6.40 | 6.16 | 0.005 | 6.10 | 5.89 | 0.004 |
| LOC100506538<br>/// NDUFAF6 | 8.13 | 7.57 | 0.001 | 8.50 | 8.28 | 0.004 | 8.11 | 7.69 | 0.008 |
| LOC100507534 | 6.38 | 5.28 | 0.002 | 6.29 | 5.51 | <0.001 | 4.42 | 4.19 | 0.007 |

These genes were classified into six categories on the basis of their molecular functions (Fig. 1). Further; the genes were classified into fourteen categories on the basis of involvement in biological processes and events (Table 2). However, some genes belong to more than one category.

**Fig. 1.**
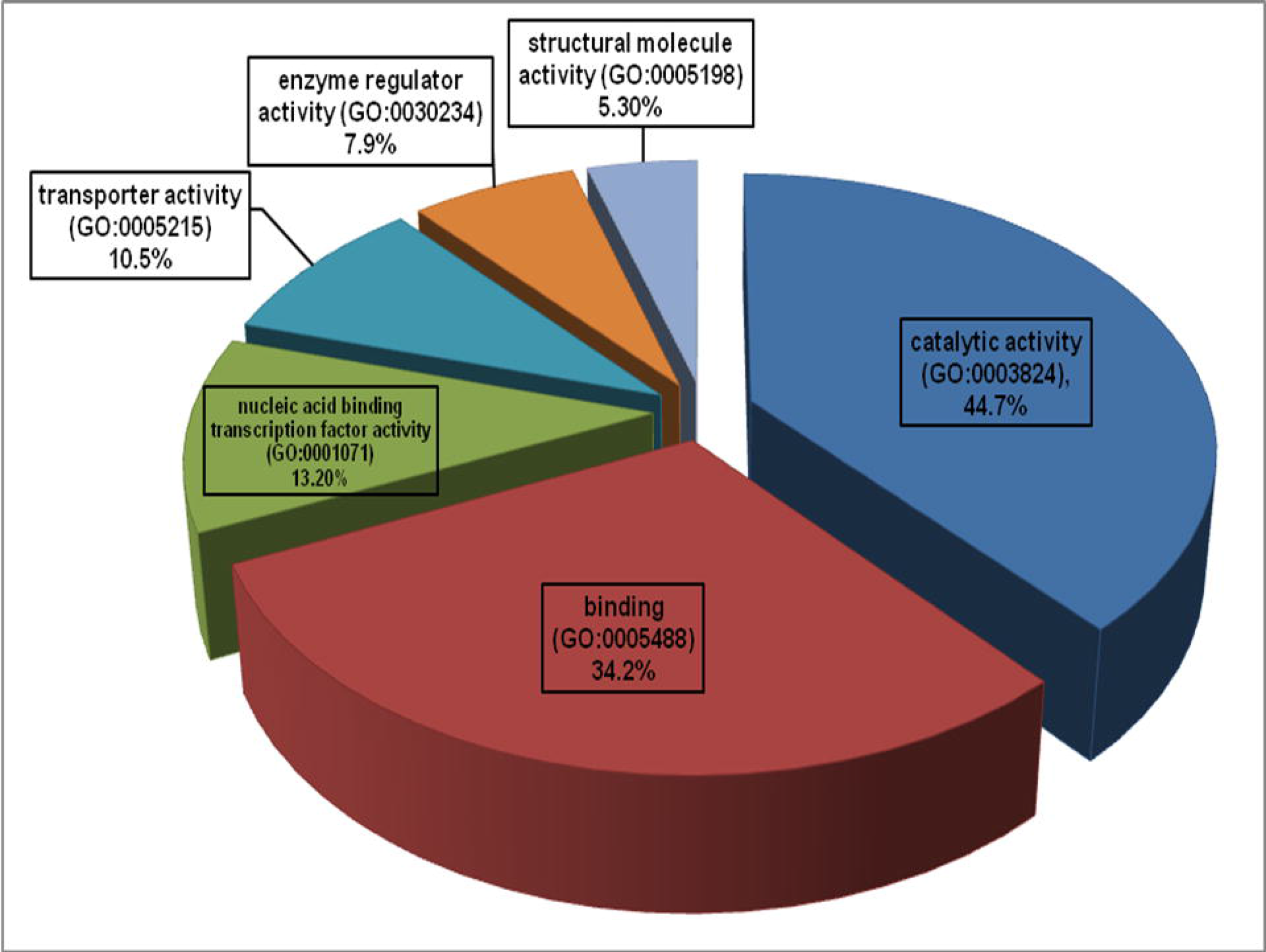
Involvement of the gene set in molecular functions. Catalysis (n=17, p=44.7%), binding (n=13, p=34.2%), and nucleic acid binding transcription factor activity (n=5, p=13.2%), transporter activity (n=4, p=10.50%), enzyme regulation (n=3, p=7.90%), and structural molecule activity (n=2, p=5.3%), n=number of genes, p=percentage Source: Panther Classification System) **Cross-references in Text**: Result and Discussion (subsections ‘Involvement of the gene set in molecular functions’ and ‘Genetic basis of schizophrenia: indications from this study’).

**Table 2.**
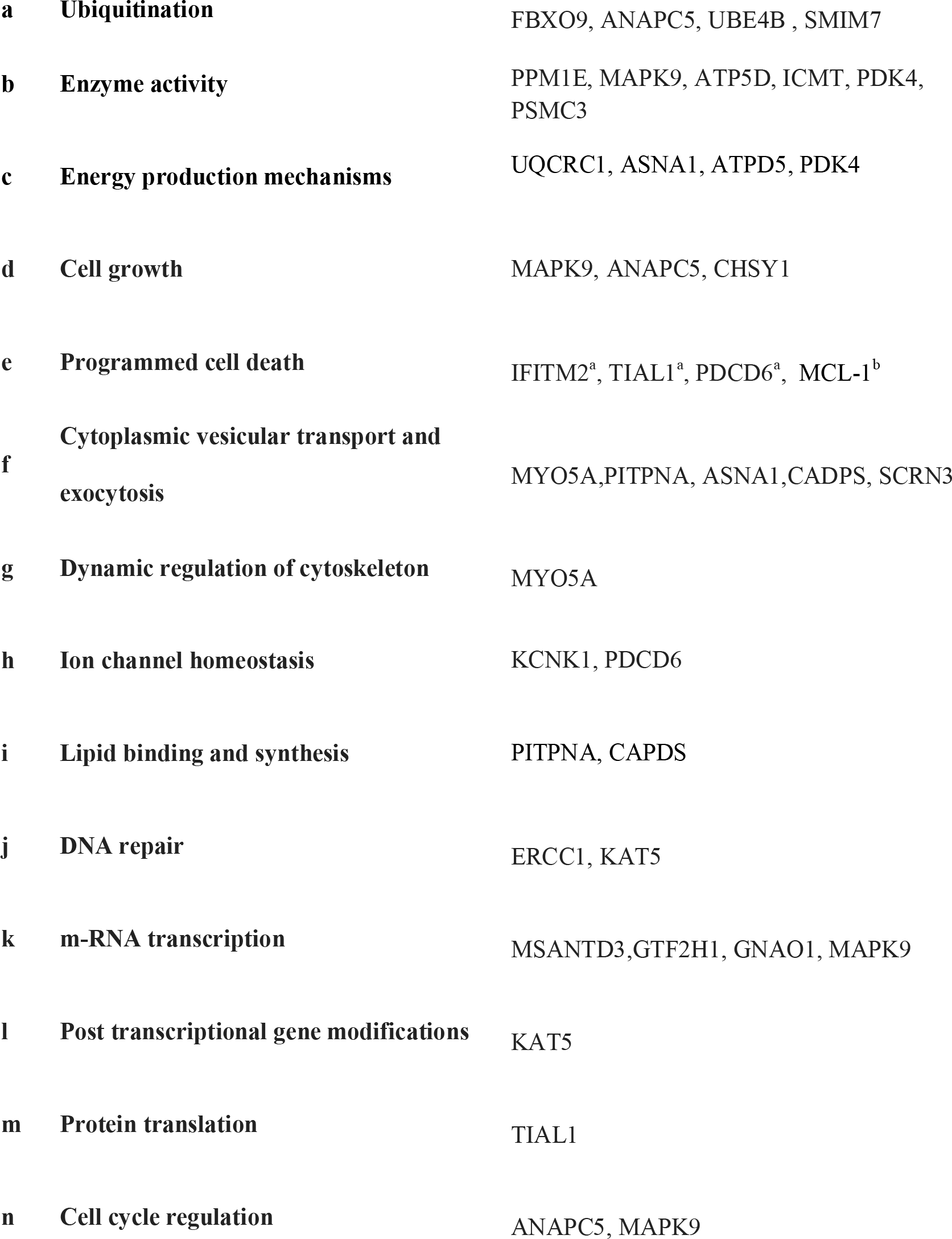
Involvement of the gene set in biological processes and events.

Reference: NCBI gene database (http://www.ncbi.nlm.nih.gov/gene/). a=Pro-apoptotic, b=Anti-apoptotic

Furthermore, in pathway linkage analysis, the gene set was found to link with 36 molecular pathways (Fig. 2) that broadly could be placed in seven categories based on their commonality (Table 3).

**Fig. 2.**
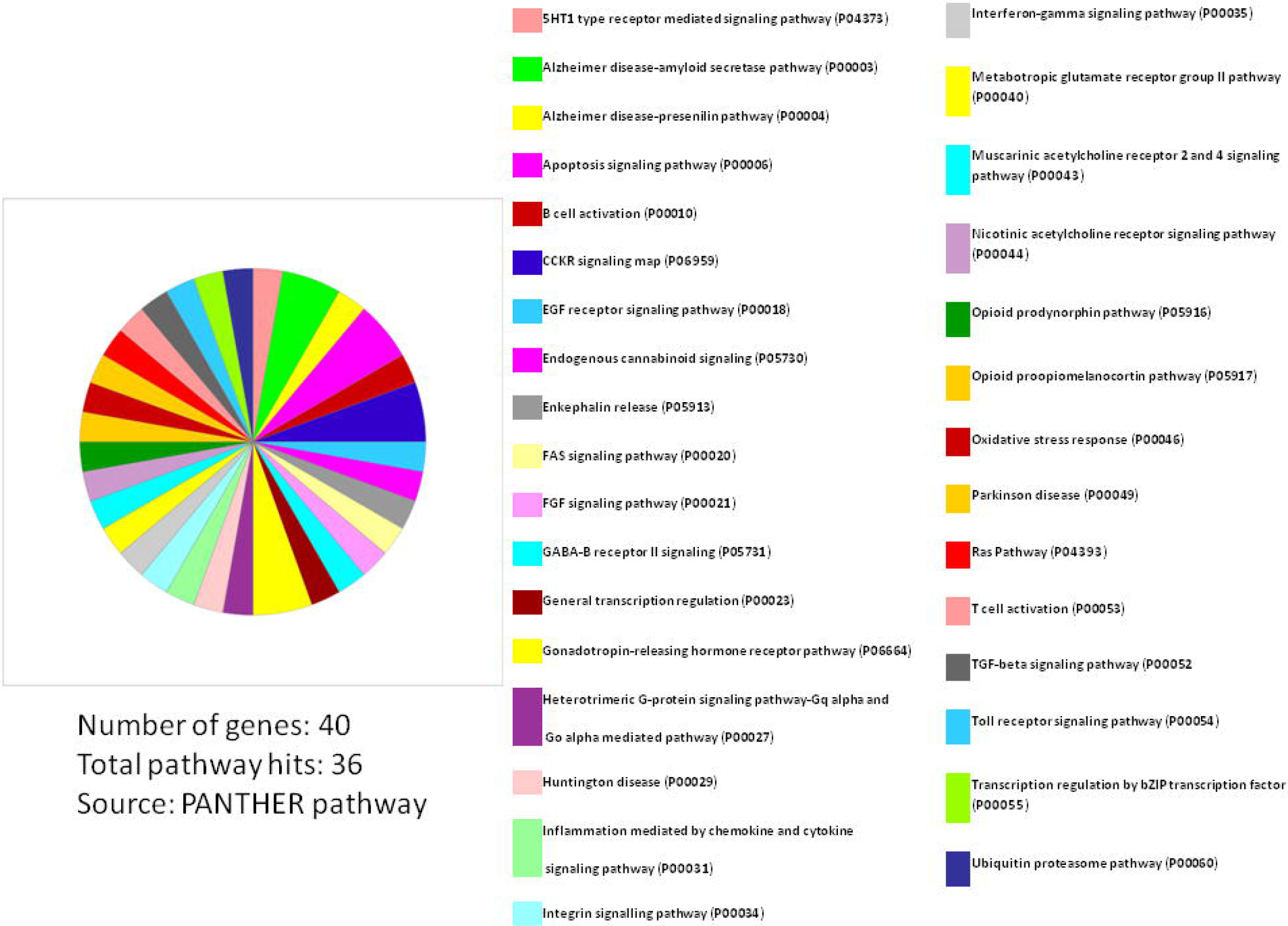
Involvement of the gene set in molecular pathways. **Cross-references in Text**: Result and Discussion (subsections ‘Involvement of the gene set in molecular pathways’ and ‘SCZ may have a neurodevelopmental and neurodegenerative component together’).

**Table 3.**
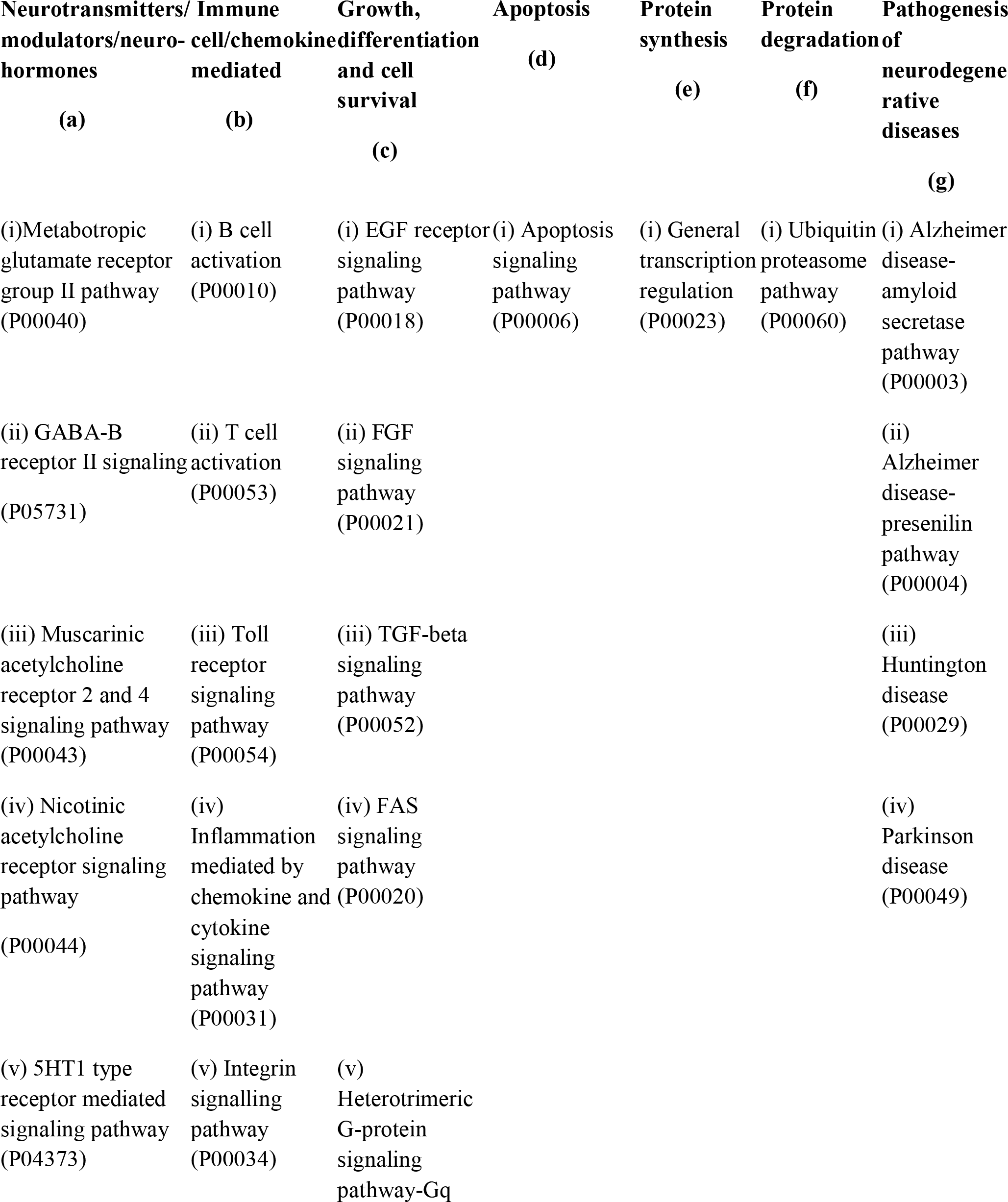
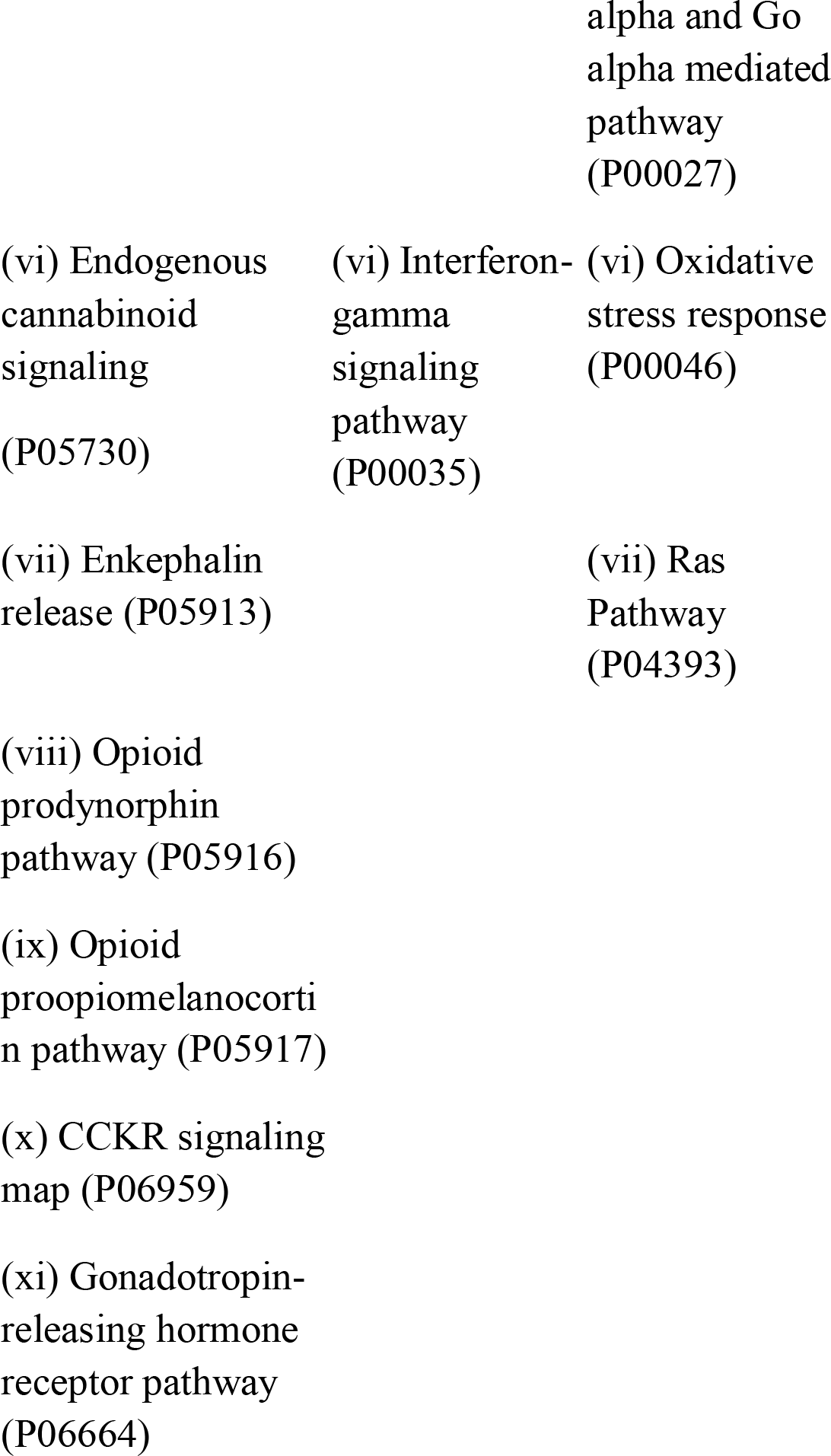
Involvement of the gene set in molecular pathways.

## Discussion

The structural and functional brain abnormalities have been repeatedly reported in patients with SCZ [65]. The brain regions chosen for this study are noted to be predominantly affected in SCZ [6-8]. Hence, the study of genome expression status in these brain regions was expected to unravel mysterious disease etiology which may be conclusive in deciding suitable therapeutic strategies to the disease. Also, as none of the genes revealed in this study had been reported earlier as the candidate gene, the new set appeals for fresh attention for the etiology of SCZ.

### Involvement of the gene set in molecular functions

The functional analysis of the gene-set (Fig. 1) elucidated the genes being involved in the regulation of basic machinery and housekeeping functions of the neurons viz. receptor-ligand binding [66], catalysis [67], enzymatic regulation [68], nucleic acid binding transcription factor activity [69], structural molecule activity [70] and transport activities [71]. It is well evident that dysregulation of all these basic functions of neurons will certainly manifest in compromised neuronal physiology and hence information processing which has been a hallmark of the progressed SCZ [72]. The molecular function analysis also showed the hierarchy of the functions compromised in SCZ (Fig. 1) (the catalysis and receptor-ligand binding being most affected functions), and that knowledge can be exploited in prioritizing therapeutic targets.

### Involvement of the gene set in biological processes and events

The comprehensive influence of the dysregulation of these genes in pathogenesis of SCZ gets further clarified in the analysis for the involvement in the biological processes and cellular events (Table 2). The implication of genes involved in ubiquitination (Table 2a), enzyme activity (Table 2b) and energy production mechanisms (Table 2c) may point towards a generalized failure of the basic functions in neurons; as ubiquitination is known to regulate the diverse spectrum of cellular functions [73] and same should be true for the genes encoding enzymes, especially those necessary for mitochondrial functions (ATP5D, PDK4) [74], regulating specific signalling pathways (MAPK9) [68] and involved in phosphorylation (ATP5D, PDK4) or dephosphorylation (PPM1E) [75,76]. The dysregulation of genes involved in energy production (Table 2c) also confirms prevailed view in the literature that energy production mechanisms get compromised in SCZ [77-79].

Again, down-regulation of genes which function as regulator of the cell growth mechanisms (Table 2d) explains reduced neuronal cell sizes, synaptic connection and brain volume in specific brain regions noted in schizophrenia [10,80], and significant upregulation of genes involved in the programmed death (Table 2e) may indicate pro-apoptotic mechanisms prevailing in particular brain regions in schizophrenia, and this inference also gets supported by some earlier studies [81-83] but that may not be a generalised feature in SCZ, as we also noted an evidence contrary to the claim also that an anti-apoptotic gene MCL-1 [84] was found significantly upregulated in all three brain regions, but there is literature evidence that although MCL-1 is an anti-apoptotic gene, it regulates cell cycle negatively hence limiting the mitosis [85].

Also, the significantly altered expression of the genes involved in cytoplasmic vesicular transport and exocytosis (Table 2f), dynamic regulation of actin and tubulin cytoskeleton (Table 2g), and ion channel homeostasis (Table 2h) [86], lipid-binding (PITPNA) and synthesis (CADPS) (Table 2i) may hint of compromised neuronal information processing in SCZ.

### Involvement of the gene set in molecular pathways

In pathway linkage analysis (Fig. 2, Table 3), the category involving largest number of molecular pathways has been that of neurotransmitters /modulators and neurohormones (Table 3a) which fits with clinical manifestations of the disease and also gets support from existing theories that the etiology of SCZ majorly may be based on dysregulation of this category of molecules [87,88].

Various neurotransmitters based hypotheses have been proposed for the etiology of SCZ [88-92] but none of them are primarily explaining causality of the diseases. The result of this study (Fig. 3, Table 3) indicates that disease etiology is not implicating any single transmitter but many of them together, warranting for an integrative study aimed at an unifying mechanism for involvement of more than one neurotransmitter and or modulators at a time [93]. The misexpression of the genes for neurotransmitters and modulators may have the greatest impact on the synaptic transmission [94,95] and oscillation coupling of the neural wave bands [96-98] hence consequently may compromise neural communications severely.

The linkage of the immune cell/chemokine mediated pathways (Table 3b) is strongly supported by literature [99-101]. An immunogenic basis of SCZ etiogenesis had also been brought forward [11,102] although counter to this hypothesis has also been placed which limits the role of immune function related genes as a solo or major factor in SCZ etiology [103]. Also, the involvement of growth, differentiation and survival of neurons in the specific brain regions (Table 3c) [104,105] (also discussed in subsection ‘Involvement of the gene set in biological processes and events’) and pathways related to apoptosis (Table 3d) [81,82,106] (also discussed in subsection ‘Involvement of the gene set in biological processes and events’), and related to protein synthesis (Table 3e) and degradation (Table 3f) [107,108] has been well documented in the literature (also discussed in subsection ‘Involvement of the gene set in biological processes and events’). The linkage of FGF signalling pathway (Table 3c) under neuronal growth, differentiation and survival to SCZ etiology has been corroborated by a freshly published study by Narla et al (2017) who regarded it as a central pathway commanding all other pathways in developing brain strengthening the view that SCZ has a neurodevelopmental etiology [109].

The linking of the pathways involved in the pathogenesis of major neurodegenerative diseases (Table 3g) such as Alzheimer [110,111], Parkinson [112], and Huntington's disease [113,114] indicates neurodegenerative nature of SCZ and may help in understanding the disease pathogenesis as well as developing new drug targets.

### Non-protein-coding genes: Unknown functions

The neuronal functions associated with 2 non-coding genes (LOC100507534, LOC100507534) couldn’t be ascertained from the literature but it's interesting to find the significant alteration of these long non-coding RNAs in SCZ which has never been reported before. There are now strong indications that non-coding genes are implicated in SCZ pathology [115,116].

### SCZ may have a neurodevelopmental and neurodegenerative component together

This had been a long time puzzle [117] that schizophrenia is a pure neurodevelopmental [83,109] disease or neurodegenerative disorder [118,119]. Although, support for neurodevelopmental etiology is getting upper hand with fresh research (also discussed in subsection ‘Involvement of the gene set in molecular pathways ‘) [109], evidence has been presented in favour of both [120]. Our study hints for a mixed etiology with involvement of molecular pathways related to brain development and neurogenesis (Table 3c) and neurodegeneration (Table 3g). A mixed etiology has been advocated by some other authors also [119,121].

## Further insights

### Massive protein derangement

The set revealed in this study mostly contained protein-coding genes (38 out of 40), and their significantly altered expression provides a clue for massive derangement of the proteome in schizophrenia which has been also suggested by a proteome-based study [122]. A recent study has indicted abnormal metabolism of some proneural proteins involved in formation of synapses as a probable etiological factor in SCZ [123].

### Evidence for associated male infertility in schizophrenia

Two of the altered genes, SOX9 and SPAG7 (Table 1), are known to regulate selective germ cell development, and spermatogenesis which may be an explanation for the associated infertility in male SCZ patients [124,125]. Also, the implication of gonadotrophin releasing hormone (GNRH) pathway (Table 3a) with the gene set provides a reason for the associated infertility in SCZ [126]. The noted linking of the schizophrenia with sex-selective genes, and also to the fertility regulating pathway are unique findings, to our knowledge never noticed before. Although abnormal response to exogenous GNRH administration is known in acute SCZ, however, an elevated secretion of prolactin which is a GNRH suppressor has also been noted in SCZ which may be a usual side effect of many anti-psychotic drugs [127].

### Genetic basis of schizophrenia: indications from this study

Our study confirms the view that SCZ etiology is polygenic and multifactorial [33], and also identifies the probable contributory factors (Fig. 1, Table 2). Although, this study has implicated the genes representing many ontological categories as we discussed above, neither of the category has been large enough to contain more than few genes, which suggests a multifactorial etiology of SCZ where all factors will add some effects rather than being a solo actor. Further, how the change in genomic architecture is translated in to the behavioural features of SCZ can be explained by the altered expression of the genes related to specific neuronal functions (further elaborated in subsection ‘Detuning of the normal neuronal gene expression may be etiomechanism of SCZ’). Role of epigenetics in etiology of SCZ [36,128,129] gets a small hint in our study also (histone acetyltransferase gene KAT5, NCBI Gene ID: 10524, Table 2l).

### Detuning of the normal neuronal gene expression may be etiomechanism of SCZ

Massive gene expression changes in three important regions of the brain which regulate different cognitive and stereotype functions [6-8] indicate that SCZ may be arising from interactions of all such individual genetic changes hence consequent accumulated effect on neural functions of the affected brain regions. As we discussed above, the ontological analysis of the gene set in our study provides the basis for comprehensive loss of neural functions in SCZ. An altered neural functionary of these brain regions which are nodal points for the neural network involved in neurocognitive functions plausibly may manifest in SCZ like symptoms. It seems that normalized expressions of these genes are necessary for optimum neural functioning and detuning of their expressions is reflected in characteristic disorganization of the behavior marked by the disease. Further, how detuning of the expression of neuronal genes lead to change of behavior may implicate the dysregulation of the usually ongoing synaptic and other neuroplastic changes in the brain. If we elaborate this theory further, it also hints that schizophrenia pathology can be reverted to the unmeasured extent with plausible normalization of the altered expression of the genes with rehabilitative or therapeutic approaches.

## Conclusions

The gene set revealed in this study provides a better representation of the brain pathology developing in schizophrenia as it included major brain regions affected in the disease hence can be used as a genetic signature for diagnosing and monitoring the disease progression as well as therapeutic effects of the drugs, and also benefits of rehabilitative practices.

The misexpression of spermatogenesis related genes is indicating a possible reason behind predominant male infertility in SCZ.

## Limitations of this study

Although the study implied a robust design to keep away the biases by randomized sampling of the test and control, a more personalised study taking more care of possible confounding factors as age, sex, progression of disease and drug intake history of each sample etc., which will also require a much larger sample size, and also the validation of the analysed data with more than one gene expression analysis techniques and methods may further augment the value of this research. Also, the limitations with the post-mortem brain samples for the gene expression analysis should be kept in mind while making inference from the findings and interpretations of this study.

## Further research

The genes which were significantly altered in only one or two and not in all three brain regions selected for study have not been included in analysis and they might carry some value in disease etiology hence should be studied individually. Moreover, the gene expression changes in many other brain regions which are known to be implicated in SCZ, needs to be investigated. A neural circuit specific analysis of the changes in gene expressions targeted to the individual neurocognitive domains may further augment the etiological clarity on SCZ. The mis-expression of spermatogenesis related genes noted in our study is a unique finding needs to be further investigated.

## Conflict of Interest

None

## Acknowledgement

We are thankful to Thomas A Lanz, contributor of the GSE53987 dataset at Gene Expression Omnibus, NCBI.

